# Managing African Swine Fever: Assessing the Potential of Camera Traps in Monitoring Wild Boar Occupancy Trends in Infected and Noninfected Zones, Using Spatio-temporal Statistical Models

**DOI:** 10.1101/2021.06.14.448288

**Authors:** Martijn Bollen, Thomas Neyens, Maxime Fajgenblat, Valérie De Waele, Alain Licoppe, Benoît Manet, Jim Casaer, Natalie Beenaerts

## Abstract

The recent spreading of African swine fever (ASF) over the Eurasian continent has been acknowledged as a serious economic threat for the pork industry. Consequently, an extensive body of research focuses on the epidemiology and control of ASF. Nevertheless, little information is available on the combined effect of ASF and ASF-related control measures on wild boar (*Sus scrofa*) population abundances. This is crucial information given the role of the remaining wild boar that act as an important reservoir of the disease. Given the high potential of camera traps as a non-invasive method for ungulate trend estimation, we assess the effectiveness of ASF control measures using a camera trap network. In this study, we focus on a major ASF outbreak in 2018-2020 in the South of Belgium. This outbreak elicited a strong management response, both in terms of fencing off a large infected zone as well as an intensive culling regime. We apply a Bayesian multi-season site-occupancy model to wild boar detection-nondetection data. Our results show that (1) occupancy rates at the onset of our monitoring period reflect the ASF infection status; (2) ASF-induced mortality and culling efforts jointly lead to decreased occupancy over time; and (3) the estimated mean total extinction rate ranges between 22.44% and 91.35%, depending on the ASF infection status. Together, these results confirm the effectiveness of ASF-control measures implemented in Wallonia (Belgium), which has regained its disease-free status in December 2020, as well as the usefulness of a camera trap network to monitor these effects.

## 1 Introduction

African swine fever (ASF), a viral disease that causes high mortality among domestic pigs (*Sus scrofa domesticus*) and wild boar (*Sus scrofa*), originates from East Africa and is regarded as one of the most important threats to the European pig industry. Recently, ASF has been re-introduced to the wild boar populations on the European mainland, presumably due to infected meat spills in the environment (1). Most likely, this spillage mediated the recent spread of ASF through a new epidemiological cycle, designated the wild boar-habitat cycle, which involves both direct and indirect viral transmissions. Direct transmissions occur through contacts among wild boar, whereas indirect cases result from viral reservoirs in the environment, such as ASF-infected carcasses (2). This new role of wild boar in the epidemiology of ASF has led to new management guidelines of wild boar populations in infected areas (3, 4). Management strategies include continuous carcass removal from the infected zone, coupled with intense culling of wild boar within a buffer zone (5). Together, these strategies are expected to effectively reduce ASF transmission by removing viral sources from the environment in the infected zone and by depleting the susceptible wild boar population in the buffer zone. The latter is essential, since the number of individuals remaining in the host population of the buffer zone will determine the probability of the spread to a noninfected area, *i*.*e*., host threshold density. In the infected zone, after the epidemic phase, culling of the remaining wild boar will determine the probability for the disease to become endemic, *i*.*e*., critical community size (6). To evaluate measures aimed at counteracting ASF, sound information on the joint effect of the disease and culling efforts on population trends of wild boar within the managed areas is crucial (3).

Over the last decade, the use of remote cameras, henceforth referred to as camera traps (CTs), has become popular when monitoring trends in medium-size to large-size mammals, including wild boar (7, 8). Photographic captures (*i*.*e*., detections) by CTs can be translated into information on the distribution and density of a focal species. However, density estimation by CTs is still hindered by imperfect detection in many cases (*i*.*e*., not detecting a focal species, when present) (9). Given the elusiveness and nocturnality of wild boar, it is among the species subjected to severely limited detectability. Moreover, traditional density estimation methodology requires individual identification, hence cannot be applied to many common species that lack natural markings, including wild boar (10, 11). Statistical frameworks such as the random encounter model (REM) allow for density estimation of unmarked populations, while accounting for imperfect detection, using CTs (12, 13). However, the need for auxiliary data collection, restricts the use of REM in many cases (14). Occupancy models on the other hand overcome both imperfect detectability and the need for individual recognition of animals, without requiring additional information. They proceed by simultaneously estimating site-occupancy and the probability of detecting a focal species, given its presence (15). Extending occupancy models to so-called multi-season site-occupancy (MSO) models, enables estimating rates expressing population changes through time. One of these rates, the extinction rate, is of prime interest when assessing the combined effect of a viral disease, such as ASF, and culling efforts on a host population.

In the current study we evaluate the use and effectiveness of a CT network to monitor wild boar population trends throughout the recent ASF epidemic in Wallonia (Belgium). The first cases from this outbreak were reported on September 13, 2018 (16). To our knowledge, only one study so far attempted to quantify the effects of ASF on a wild boar population using CTs (17). Here, we develop a different statistical framework, that can be adapted to model management strategies of multiple species, in diverse settings. We apply it in our study of wild boar population dynamics in an ASF-infected and noninfected zone. The latter has been subjected to an intensive culling regime (January 2019 – March 2021) during the entire ASF episode. While in the infected zone, both ASF-induced mortality and culling efforts determine wild boar extinction rates. Moreover, this zone is fenced off from the surrounding noninfected zones. Using detection-nondetection data (March 2019 - May 2020) from 92 CTs, we estimate monthly site-occupancy of wild boar in both ASF-infected and noninfected zones, through a Bayesian MSO. In addition, we provide wild boar distribution maps for the area under study. Finally, we draw conclusions about the ASF management in Wallonia (Belgium) and provide recommendations regarding study design and statistical analysis for monitoring future outbreaks. We believe this case is important beyond its own setting for two reasons: (i) wildlife managers are often asked by funders to justify the use of CT networks, hence being able to show the effectiveness of CTs to monitor population trends is important; (ii) authorities need to assess the effectiveness of control measures taken to prevent the spreading of ASF. Camera trapping is an understudied method that could provide valuable information on the effectiveness of these measures.

## 2 Materials and Methods

### 2.1 Study area

The study area (longitudes: 5.2847°E - 5.8156°E; latitudes: 49.5485°N - 49.6955°N) is situated between the cities of Virton (South), Florenville (Northwest) and Arlon (Northeast), and has been subdivided into three management zones. An ASF-infected zone (red), a noninfected zone, *i*.*e*., free of ASF (turquoise) and a zone that was excluded from the study due to its ambiguous disease status and limited number of deployments (grey), see **Figure1**. It has a total forested area of 223 km^2^, part of a larger ASF management zone of 1106 km^2^ encompassing a total of 572 forested km^2^. The study area has a cool temperate and moist climate, with a mean annual temperature of 8.94°C and 966 mm rainfall (18). The landscape is characterized by rugged terrain crossed by numerous rivers and fragmented by several roads, while its vegetation is dominated by deciduous forests.

**Figure 1:**
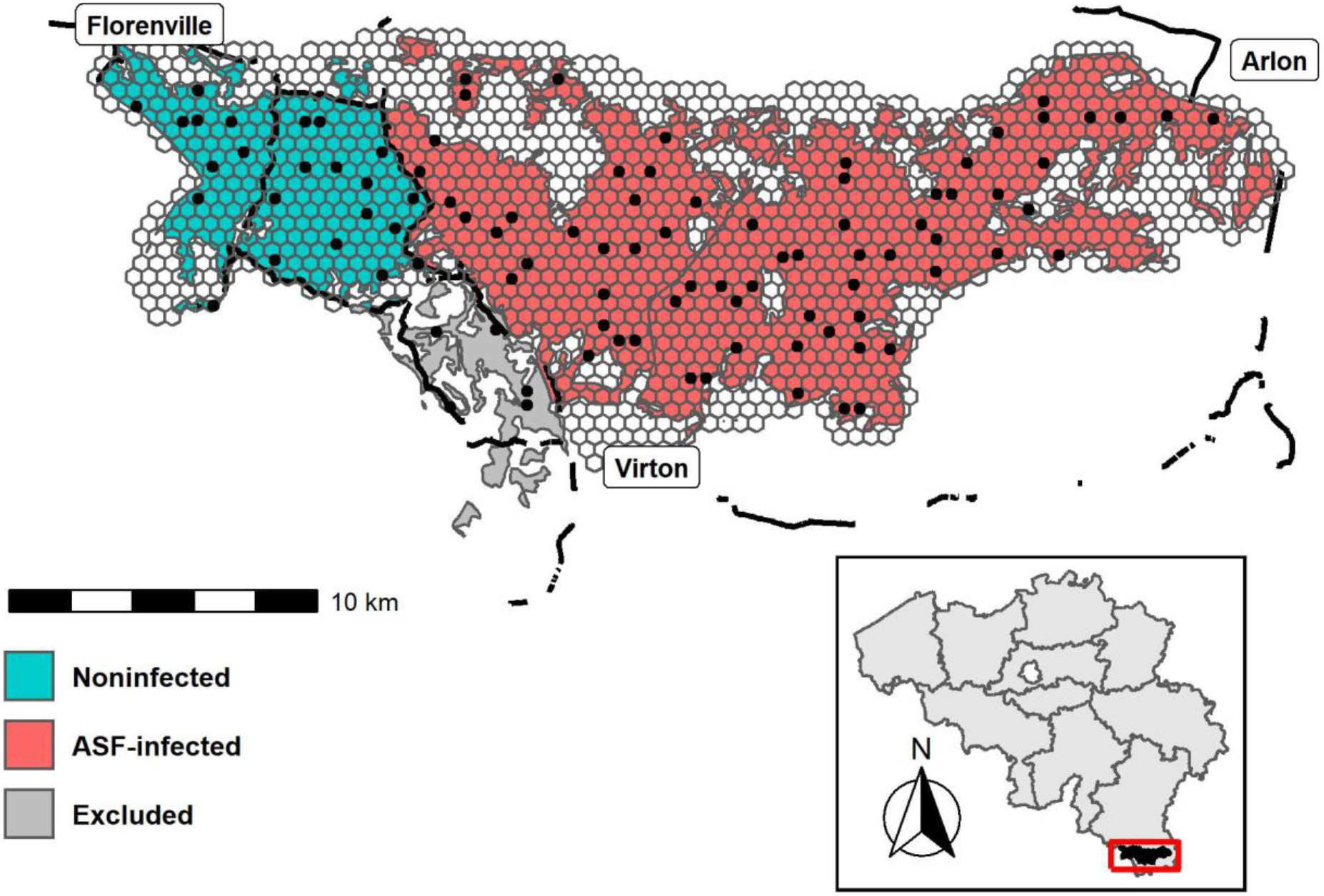
Map of the study area with the overlaying hexagonal grids. Camera deployments are indicated by the black dots. Black lines represent fences. Colors represent the African swine fever management zones; ASF-infected (red), noninfected (turquoise) and excluded (grey). The inset map (bottom) shows the study area within Belgium.

### 2.2 Camera trap network and data

Within the study area, a CT network was deployed since March 2019. The network consists out of 97 Snapshot Extra Black 12.0 l HD (Dörr) cameras. For more detailed information on the camera specifications, consult **Supplementary Table 1**.

Camera placement was done according to a stratified random sampling scheme. Proportional to its area, a number of cameras was deployed in each management zone (stratum) (**Supplementary Table 2**). *A posteriori*, the strata were superimposed by a hexagonal grid layer (x-spacing of 500 m, area of 21.65 ha/ site) ensuring that each camera was assigned a unique grid cell (**Figure 1**). Throughout the sampling period, camera locations were fixed. All cameras were installed by mounting them on trees approximately 50 centimetres above ground, facing North. We did not use baiting, nor did we select for trails. Monthly check-ups were performed to determine battery levels and to verify camera operability. Each camera trigger was followed by a series of five photographs, without a delay between consecutive triggers. All images were manually annotated, using the Agouti software platform (19). After omitting data from the excluded zone (5 deployments; 5.15%) (**Figure 1**), we retained data from 92 deployments between March 2019 and May 2020, resulting in a total of 42,136 24-h observation periods.

We considered three classes of covariates, potentially important to explain wild boar population trends: (1) time, (2) land use, and (3) infection status. As will be clarified in Section 2.3, for time, we evaluated three alternative definitions: (1.i) month since the start of the monitoring program (observation month; *t*), (1.ii) a binary variable indicating whether the observation month is in April – September (biannual;), and (1.iii) a similar indicator variable consisting of four seasonal periods, *i*.*e*., Spring, Summer, Autumn and Winter (quarterly; *QRT*_*t*_). For land use, we only considered a single covariate: the (*z*-scored) proportion of broad-leaved forest land cover class (*BL*_*t*_), which was extracted from the LifeWatch Ecotope dataset (18). Finally, infection status (*ASF*_*i*_) is encoded as follows: 1 when a site is within the ASF-infected zone, 0 otherwise (**Figure 1**). An overview of these covariates and *a priori* defined models are given in **Table 1**. For model-specific predictions (P(*Model*)) consult **Supplementary Table 3**.

**Table 1:**
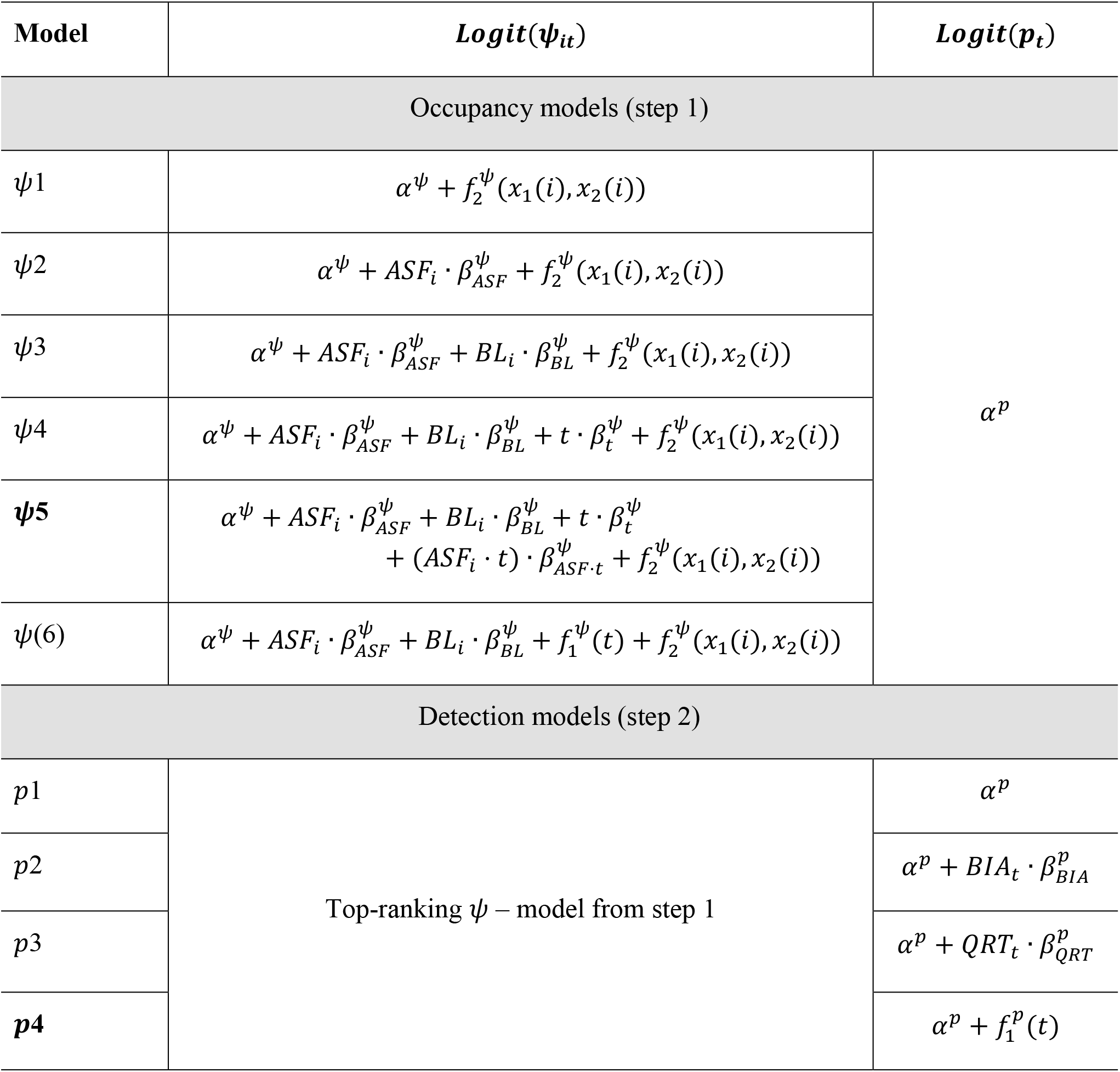
*A priori* defined occupancy (step 1) and detection (step 2) models. Top-ranked models for each step are indicated in bold. Abbreviations: ASF infection status (*ASF*_*i*_), *z*-scored proportion of broad-leaved tree land cover class (*BL*_*i*_), observation month (*t*), biannual (spring – summer, autumn – winter) seasons (*BIA*_*t*_), quarterly (spring, summer, autumn, winter) seasons (*QRT*_*t*_), smooth function for temporal variation *f*_1_(*t*), smooth function for spatial variation *f*_2_(*t*). Intercepts and slope parameters are given by *α* and *β* respectively.

### 2.3 Statistical model

We analyse the CT data using a longitudinal multi-season occupancy model (MSO), defined as a state-space model (20), to make inference on wild boar’s site-occupancy. The sampling grids used are smaller than wild boar’s home range, hence occupancy should be interpreted as habitat use (21). Detection histories were constructed using the R package *CamtrapR* (22). For site *i* = 1,2, …, *N*, at survey day *j* = 1,2, …, *J*, in observation month *t* = 1,2, …, *T*, the detection history is 1 when wild boar were observed during a 24-h period (*y*_*ijt*_ = 1) or 0 when no boars are caught on camera that day (*y*_*ijt*_ = 0). These are assumed to follow a Bernoulli distribution, such that,

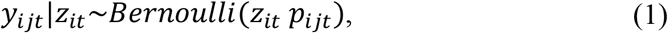

where *p*_*ijt*_ is the probability of detecting the focal species and *z*_*it*_ is the latent occupancy status (unoccupied *z*_*it*_ = 0; occupied *z*_*it*_ = 1) at site during observation month *t*. Note that we do not use survey day-specific, nor site-specific covariates to model *p*_*ijt*_, hence the detection probability simplifies to *p*_*t*_. The occupancy status is modelled as,

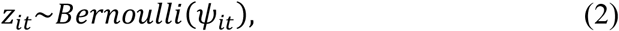

Where *Ψ*_*it*_ is the occupancy probability, from now on simply referred to as ‘occupancy’, at site during observation month *t*. Unlike dynamic MSO, we do not take probabilities of wild boar surviving or colonizing a site *i* from observation month *t* to *t* + 1 into account, as the high degree of zero-inflation in our data complicates joint inference on all these processes. We define 𝒱_*l*_ = {*pijt, Ψ*_*it*_}, which collects all processes that will be modelled as a function of covariates and random effects, using a logit link. A general model formulation for 𝒱_*l*_, *l* = 1, 2, can be defined as

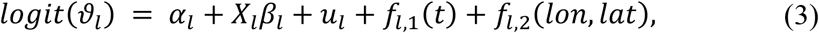

where *α*_*l*_ are intercepts, *β*_*l*_ are vectors of process-specific slope parameters with their corresponding covariate matrix *X*_*l*_. The term *u*_*l*_ models spatially unstructured overdispersion as a normally distributed random effect, *f*_*l*,1_ is a smooth function modelling temporal variation for each observation month *t* and *f*_*l*,2_ is an isotropic two-dimensional smooth function modelling spatial variation in occupancy patterns, for the longitude *lon* and latitude *lat* of each site’s centroid. Both *f*_*l*,1_ and *f*_*l*,2_ are modelled using Gaussian processes (GP), with an exponentiated quadratic covariance function. While *f*_*l*,1_ uses an exact GP, we model *f*_*l*,2_ by means of the Hilbert space reduced-rank Gaussian process (HSGP) approach as the number of sites in our study area is large (23, 24). This approach yields substantial speed gains when dealing with large number of sites through approximate series expansions of the GP’s covariance function.

Model fitting was performed using *Stan* (via the R package *rstan*), a probabilistic programming language that enables Bayesian estimation through a dynamic Hamiltonian Monte Carlo (HMC) sampler (25). For each MCMC iteration, we also derive site-specific growth rates 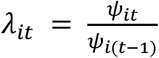, average monthly growth rates 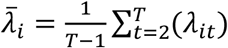 and total growth rates 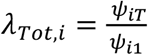 in site-occupancy (26). We choose weakly informative *Student t*(3,0,5) priors for all the regression parameters {*α*_*l*_, *β*_*l*_}and a nonnegative *Student t*^+^ (3,0, 5) prior for the marginal standard deviation of the hyperparameters 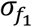 and 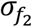 of the GPs. For the scale parameters 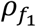 and 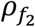 of the GPs, we respectively used an inverse gamma *IG*(10.9, 4.00) and a generalized inverse Gaussian *GIG*(3,12,0.01), ensuring most prior evidence is placed on scales that can be estimated from the data (*i*.*e*., larger than the smallest difference between any pair of CT locations and smaller than the largest difference between any of these pairs).

The full model that would contain two random effects terms for each of these processes, in addition to covariates, was computationally infeasible to fit and, furthermore, does not necessarily reflect a sensible data-generating process. Hence, we consider a set of sensible reduced models based on ecologically plausible considerations, preventing multicollinearity, and computational feasibility (**Table 1**). Multicollinearity was avoided by including one of two covariates, when their Spearman rho correlation estimate |*r*_*s*_| < 0.6. Subsequently, we select the most appropriate model by means of a model selection procedure.

Model selection through approximate leave-one-out cross-validation was performed using the R package *loo* (27). Following the authors’ recommendations, leave-one-out (LOO) expected log-predictive densities were used to rank our *a priori* selected candidate models (**Table 2**). Our ranking procedure consists of a two-step approach. First, the top-ranked occupancy model is retained by comparing LOO for selected combinations of fixed and random effects at the occupancy (*Ψ*) level, while keeping detectability (*p*) constant (**Table 2**, step 1). Secondly, the detection process is modelled using fixed effects only, while adopting the top-ranked occupancy model from step 1 (**Table 2**, step 2).

**Table 2:** Model selection of candidate occupancy models (step 1) and detection models (step 2). Note that P_D_ represents the effective number of parameters rather than the true number of parameters. Leave-one-out expected log-predictive density (LOO), in addition to the difference in LOO between each model and the top-ranked model (Δ LOO), along with their standard errors (SE) are presented.

| Model | $P_D$ | SE($P_D$) | LOO | SE(LOO) | $\Delta$ LOO | SE($\Delta$ LOO) |
| --- | --- | --- | --- | --- | --- | --- |
| Occupancy models (step 1) |  |  |  |  |  |  |
| $\psi(5)$ | 45.55 | 2.84 | -992.29 | 60.04 | 0.00 | 0.00 |
| $\psi(4)$ | 43.49 | 2.70 | -995.43 | 60.03 | -3.14 | 2.38 |
| $\psi(6)$ | 53.02 | 3.25 | -999.16 | 60.20 | -6.87 | 3.56 |
| $\psi(3)$ | 40.92 | 2.44 | -1044.70 | 61.14 | -52.41 | 9.84 |
| $\psi(2)$ | 38.64 | 2.32 | -1045.91 | 61.12 | -53.62 | 9.94 |
| $\psi(1)$ | 37.83 | 2.33 | -1049.73 | 61.24 | -57.44 | 10.45 |
| Detection models (step 2) |  |  |  |  |  |  |
| $p(4)$ | 71.17 | 6.93 | -979.88 | 59.33 | 0.00 | 0.00 |
| $p(2)$ | 47.31 | 2.96 | -985.54 | 58.15 | -5.66 | 9.41 |
| $p(1)$ | 45.55 | 2.84 | -992.29 | 60.04 | -12.40 | 10.88 |
| $p(3)$ | 47.26 | 2.92 | -993.16 | 60.21 | -13.28 | 10.49 |

All models were fitted using four parallel MCMC chains with 4000 iterations, which included 2000 iterations that were discarded as burn-in iterations for all candidate models; this always resulted in satisfactory convergence (**Supplementary Figure 1**), following the guidelines by Vehtari et al. (28). After the selection procedure, a prior sensitivity analysis was performed for the top-ranked model from step 2, by comparing results of the default prior specification with *Student t*(3, 0, 2.5) and *Student t*(3,0,10) priors for 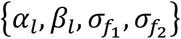 this analysis revealed posterior invariance under the considered prior specifications.

## 3 Results

**Table 2** presents the model selection process, which yielded a final model consisting of an occupancy process and detection process that will be detailed in the following subsections.

### 3.1 Detectability

The detection model according to LOO (**Table 2**, *p*4) models temporal variation in wild boar’s detectability for each observation month as a GP. Modelling the detection probability using biannual seasons (*p*2) also resulted in a better fit compared to the intercept only model (*p*1). Using quarterly instead of biannual seasons, resulted in the lowest ranking detection model (*p*3). Note that the accuracy in Δ LOO, measured as the standard error of this metric, is relatively low for all detection models (**Table 2**, step 2). The posterior mean probability of detecting wild boar ranges between 0.0279 and 0.1106 regardless of the observation month. Despite low detectability in general, monthly differences can be observed (**Supplementary Figures 2-3**).

### 3.2 Occupancy

All of the tested covariate combinations performed better than the intercept model (*Ψ*1), with the multiplicative model of *ASF*_*i*_, *t*, and *BL*_*t*_ (*Ψ*5) outranking all other models. Similar to the detection models, not all Δ LOO values are accurate. For the difference between the two top-ranking models the standard error exceeds |Δ LOO| (**Table 2**, step 1). According to the top-ranked occupancy model, ASF infection status has a significant effect on the occupancy of wild boar. Posterior mean odds ratios (OR) of wild boar occupancy are 17.71 (3.49 – 95.12) and 0.01 (0.00 – 0.08) for noninfected and ASF-infected zones respectively. Moreover, the OR for season, 0.76 (0.65 – 0.88), reveals a significant overall decline in wild boar occupancy for observation month (*t*). Finally, the OR for both proportion of broad-leaved forest land cover class (*BL*_*T*_) and the ASF infection status – season interaction term (*ASF*_*i*_. *t*) is insignificant, *i*.*e*., one is enclosed by the 95% HPDI (**Supplementary Table 4)**. Posterior mean occupancy and 95% HPDI at each observation month, averaged over the ASF management zones (**Figure 2**), reveal an overall decline in occupancy, both in the ASF-infected and noninfected zone. Finally, prediction maps for the estimated occupancy from March 2019 until May 2020, are displayed in **Figure 3** (see **Supplementary Figures 4-5** for 2.5^th^ and 95^th^ percentile maps).

**Figure 2:**
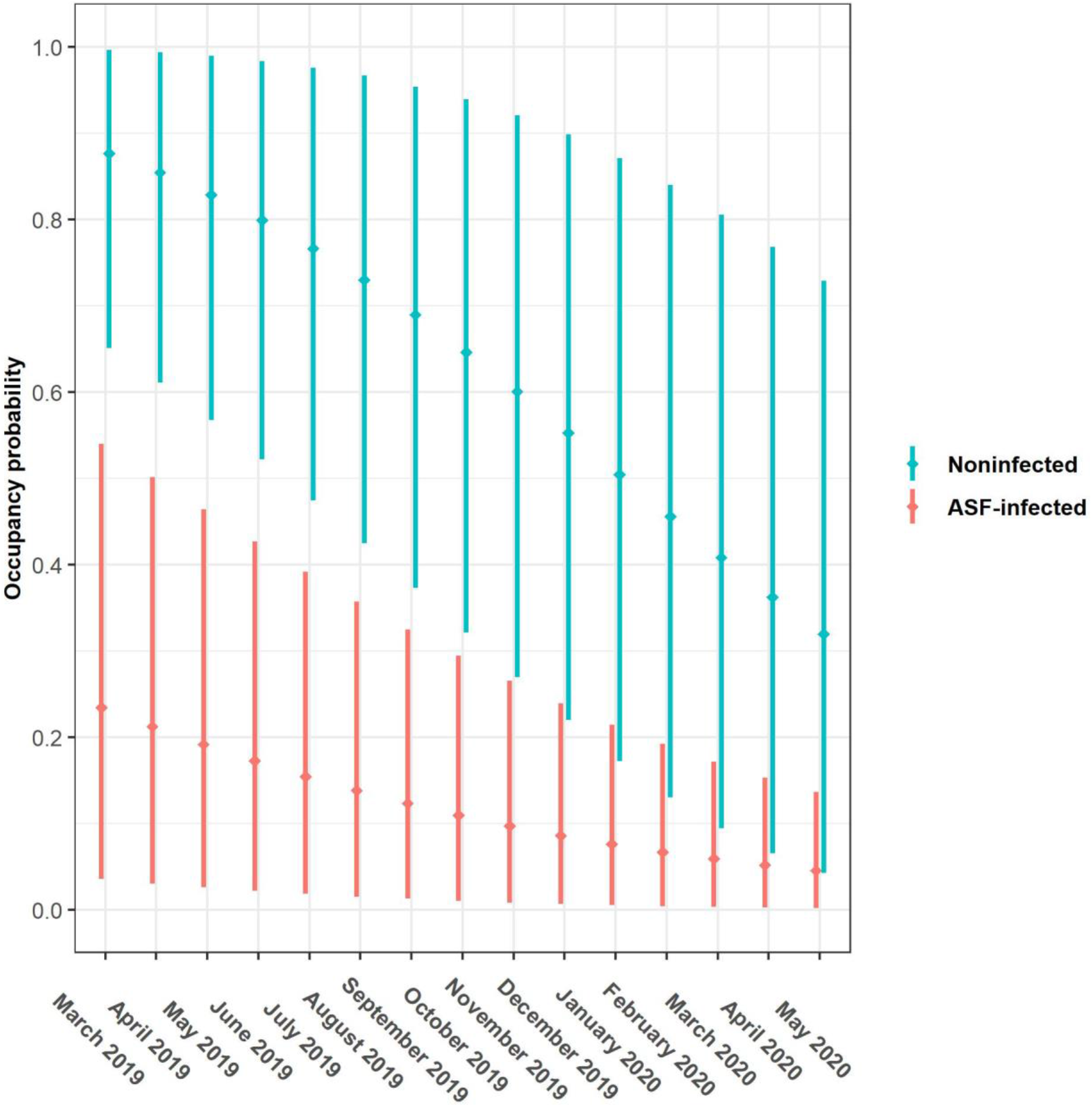
Monthly posterior mean occupancy estimates (dots) and 95% highest posterior density intervals (vertical lines) per ASF management zone.

**Figure 3:**
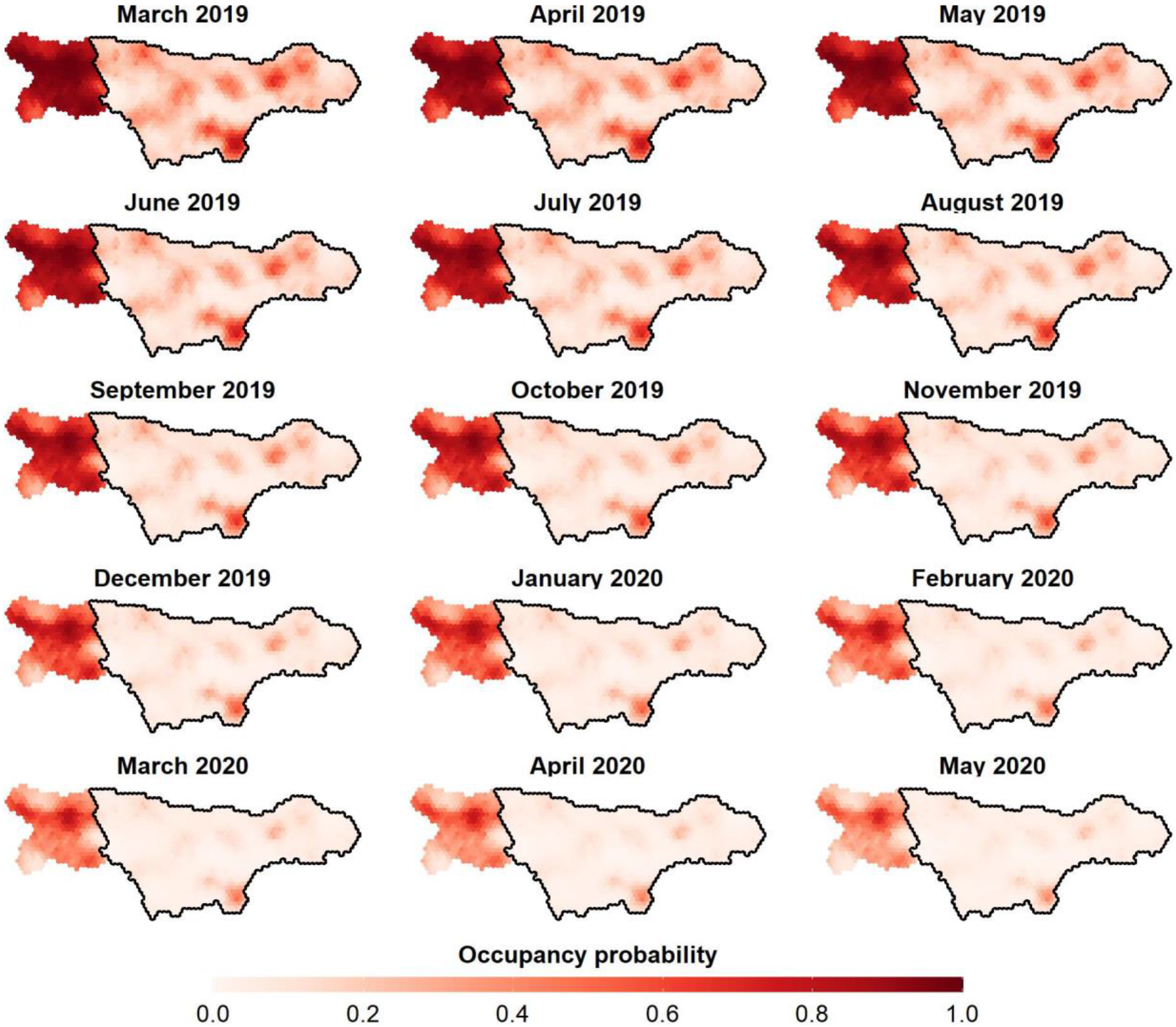
Posterior mean occupancy of wild boar in the ASF-infected (enclosed by the black line) and noninfected (non-enclosed) zone in Wallonia (Belgium). Panels ranging from March 2019 until May 2020.

### 3.3 Occupancy growth rate

Posterior means of occupancy growth rates, *i*.*e*., *λ*_*it*_, are lower than one regardless of the site and season (**Supplementary Figure 6**). For *λ*_*Tot,i*_, total growth rates (in fact, extinction rates, due to their negative trend) in occupancy, posterior means range between 0.0865 and 0.7756 (0.9135 and 0.2244), while those for average monthly growth rates 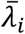 lie between 0.8049 and 0.9757 (0.1951 and 0.1243). Finally, posterior mean and 95% HPDI for *λ*_*Tot,i*_ and 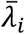 averaged over the ASF management zones (designated *λ*_*Tot,z*_ and 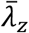) are given in **Table 3**.

**Table 3:** Posterior mean and 95% highest posterior density values for the total growth rate (*λ*_*Tot,z*_) and average monthly growth rate 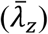 per ASF management zone, obtained by averaging over all corresponding sites.

| Zone | Mean | 2.5% | 97.5% | Mean | 2.5% | 97.5% |
| --- | --- | --- | --- | --- | --- | --- |
| | $\bar{\lambda}_z$ | | | $\lambda_{Tot,z}$ | | |
| ASF-infected | 0.8670 | 0.8083 | 0.9264 | 0.1772 | 0.0494 | 0.3432 |
| Noninfected | 0.9032 | 0.8105 | 0.9900 | 0.3848 | 0.0739 | 0.7946 |

## 4 Discussion

To assess the effectiveness of CTs to monitor wild boar population trends throughout the recent ASF epidemic in Wallonia (Belgium), we have built a spatio-temporal MSO model. This was done according to a two-step approach, selecting the best detection covariates and subsequently occupancy covariates with respect to the LOO (27) statistics from a set of *a priori* defined models.

### 4.1 Detectability

For all model comparisons (relative to the top-ranked detection model) the standard errors for Δ LOO values are smaller than two times |Δ LOO|, hence a certain degree of uncertainty as to which model provides the best fit to the data remains after our selection procedure (**Table 2**, step 2). Nevertheless, we believe that using a GP to model monthly temporal variation in wild boar detection probability is a sensible choice, given the ability of GPs to balance ecological realism with model flexibility (29).

When using CTs, detection probabilities are known to be affected by, among others, vegetation denseness, background surface temperature and weather conditions (30), all of which depend on the seasonal variation to some extent. Hence, a certain degree of seasonality in detection probabilities is not uncommon. Morelle et al. (17) report higher probabilities of detecting wild boar in summer months compared to fall (4.90E+04), winter (4.34E-03) and spring (4.90E+04). In this study, posterior mean detection probability for wild boar is low, although some additional heterogeneity attributed to the observation month was observed. In 2019, summer months display a somewhat higher probability of detecting wild boar as compared to winter months, yet there is insufficient evidence that a periodic trend exists. We rather suggest that the main effect at play, is a density-dependent effect (31, 32), more specifically a decline in detection probability that coincides with a decreasing wild boar density. In addition, the intensive culling regime adopted throughout the ASF epidemic possibly led to an increased risk perception by wild boar, incentivizing them to restrict their movements and seek hiding places. Lower activity levels negatively relate with photographic rates (33). Similarly, low probabilities of detecting wild boar could reflect decreased movement.

### 4.2 Occupancy

The top-ranked occupancy model consists of a multiplicative effect the ASF infection status (*ASF*_*i*_) of the zone, the observation month (*t*) and the proportion of deciduous forest land-use class (*BL*_*t*_), which is followed by a fully additive model of these covariates (**Table 2**, step 1). A large difference in wild boar occupancy was already present during the first month of the study period (March 2019), with posterior mean occupancies of 0.2352 and 0.8677 in infected and noninfected zones respectively (**Supplementary Table 5**). Hence, ASF-governed mortality, which had already decimated the population in the infected zone at the beginning of the monitoring program, strongly affects wild boar occupancy (posterior mean 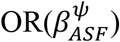 of 0.01; **Supplementary Table 1**). This is not surprisingly given that the zone was already infected with ASF for several months (*i*.*e*., since September 2019) before the study’s onset. Although it is uncertain whether the low initial occupancy in the infected zone is driven by ASF alone, mortality rates approaching 100% have been reported (34). Interestingly, the inclusion of a HSGP that accounts for extra-variability due to spatial correlation did not prevent the existence of a strong boundary between the ASF-infected and noninfected zones with respect to their occupancy patterns (**Figure 3**). This boundary effect remained, even after omitting the information about ASF infection status from our model (results not shown). In that case, the variation in occupancy previously accounted for by 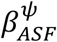, was taken up by the spatial GP 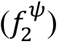. Hence, we are confident that this boundary effect is not an artefact. Instead, we argue that a sudden rise in occupancy at the infected-noninfected boundary results from having fenced off the ASF-infected zone (**Figure 1**). The latter effectively creates two separate subpopulations with each of them subjected to distinct population dynamics, leading to an unprecedented extinction rate in the infected zone.

Despite the strong impact of ASF, the infected zone seems to have had quite some refugee sites that display higher occupancies compared to surrounding areas as of March 2019 (**Figure 3**). As the epidemic progressed, occupancy dropped in most of these subareas, with only one patch in the South displaying markedly higher occupancy towards May 2020. Given the remoteness of this patch, it could be that wild boar in this area were shielded from ASF to some extent. A more likely explanation is that these refugee sites reflect the area’s suitability for remaining wild boar in terms of habitat quality and food availability. Similarly, we argue that latent ecological preferences drive the heterogeneity in occupancy observed within the noninfected zone, where higher occupancies were observed in the central axis (horizontally) throughout the study period (**Figure 3**). Indeed, looking at the spatial random effect alone, both the Southern patch of the ASF-infected zone and central axis of the noninfected zone are associated with some of the highest values (**Supplementary Figure 7**). A number of potential ecological drivers of wild boar occupancy were observed and subsequently modelled; we have considered the proportion of broad-leaved tree land cover class, which is known to positively affect its occupancy (35-37), as a fixed effect in our final model. We find this covariate to be not significant at the 5% significance level, but significant at the 10% level (results not shown), which suggests an effect that needs further investigation in future studies.

Finally, we obtain an overall declining trend (posterior mean 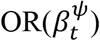 of 0.76; **Supplementary Table 4**) in wild boar occupancy for both ASF-infected an noninfected zones (**Figure 2**). This indicates that occupancy probabilities continue to drop as a consequence of the joint effect of ASF-induced mortality and culling efforts in the infected zone. While the decline in occupancy in the buffer zone results from culling alone. Together, these findings support the management strategies adopted in Wallonia (Belgium). In addition, we find that occupancy declines have different rates between the zones, with a more moderate decline seen in the ASF-infected zone (posterior mean 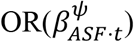 of 1.13; **Supplementary Table 4**). Possibly, differences in hunting pressure could explain this variation in rates of occupancy decline. Although wild boars have been culled in both ASF-infected and noninfected zones, the latter was more densely populated throughout the entire study period, which likely reduces the search effort by hunters and leads to increased hunting success (**Supplementary Table 6**). In addition, we note that between the epidemic-onset and its peak (September 2019 – February 2019), an occupancy decline was likely much higher in the ASF-infected zone. Importantly, our model reveals that the *ASF*_*i*_ 302 · *t* interaction term is insignificant when looking at 95% HPDI. Uncertainty about the existence of zone-specific occupancy rates, is also reflected in the small Δ LOO between a model with and one without the interaction term (**Table** 2, step 1).

### 4.3 Occupancy growth rate

We will not discuss growth rates in depth, since they carry on the same messages as the occupancy probabilities discussed earlier, but see **Supplementary Figure 6** for a graphical representation. However, it is worthwhile to briefly focus on total occupancy growth rates, as they provide a summary statistic for net change in occupancy. Posterior means of 0.1772 and 0.3848 (extinction rates of 0.9228 and 0.7152) for respectively ASF-infected and noninfected zones, confirm the strong decline in wild boar occupancy. In line with these results, Morelle et al. (17) report declines in wild boar abundance, obtained through fitting a REM (12) to CT data, of 83.8 ± 25.5% and 94.8 ± 6.4% in unmanaged and managed zones respectively, one year after an ASF outbreak in the Bialowieza Primeval Forest (Poland). Moreover, average monthly growth rates of 0.8670 and 0.9032 (**Table** 3) indicate that monitoring highly lethal diseases, such as ASF, which typically lead to rapid depletion of individuals, demands for shorter primary sampling periods.

### 4.4 Limitations

The data used in this study do not cover the full ASF episode as it occurred in Wallonia (Belgium). As such, we are unable to report on the full course of the epidemic. Secondly, it has been reported by (26) that sample sizes smaller than 40 lead to insufficient power to detect declines in occupancy under most circumstances. Here, we deploy 69 cameras in the ASF-infected and only 23 cameras in the noninfected zone. Nonetheless, we were able to capture significant declines in occupancy for both zones throughout the study period. Importantly, both ASF management zones had sampling intensities higher than the best scenario (2% of sites sampled) considered by Banner et al. (26). From a modeller’s perspective, we did not attempt a full spatio-temporal analysis. However, we believe that it is reasonable to assume that temporal dynamics in site-occupancy are spatially independent, given the relatively small surface area (ASF-infected: 162.826 km^2^, noninfected: 48.229 km^2^) of both ASF management zones. Finally, we did not include structured and unstructured random effects for both the detection and occupancy process, due to unidentifiability.

### 4.5 Conclusion

Based on our results, we conclude that ASF infection status was the main driver of wild boar occupancy at the beginning of the monitoring period, which results in a clear difference in occupancy between ASF-infected and noninfected zones. Moreover, we find that fencing off the infected zone has helped to maintain this sharp contrast in occupancy probability throughout the study period. Starting from March 2019, our model strongly supports an overall decline in occupancy until May 2020. Additionally, we attribute a steeper decline in the noninfected zone to the management strategies, adopted to counteract the progression of ASF. Together, these results confirm (1) the effectiveness of ASF control measures implemented in Wallonia (Belgium), and (2) the potential of using a CT-network to monitor the impact of both the disease and the management actions on wild boar populations during an ASF outbreak.

## 6 Author Contributions

Each author’s contribution is described using the CRediT roles.

**MB:** Methodology, Formal analysis, Visualization, Writing – Original draft preparation. **TN:** Methodology, Validation. **MF:** Methodology, Visualization. **AL:** Resources. **VDW:** Resources. **BM:** Resources. **JC:** Supervision. **NB:** Supervision. **All authors:** Writing – Review & Editing, Conceptualization (all except MF and BM).

## 8 Acknowledgments

We thank Guillaume Morrel for installing the camera trapping network during his master thesis (ULiège). Further, we are grateful to the municipalities and residents in Gaume to allow us to place camera traps on their property.

## 9 Data Availability Statement

The datasets generated for this study can be found in [figshare] [https://figshare.com/projects/African_Swine_Fever_Monitoring/115092].

## 10 Ethics statement

This work was not subject to permission/authorization from an ethical commission, since we used a non-invasive method (camera trapping) which does not disturb the natural behavior of animal.

