## Supplementary figures and tables for "Managing African Swine Fever: Assessing the Potential of Camera Traps in Monitoring Wild Boar Occupancy Trends in Infected and Noninfected Zones, Using Spatio-temporal Statistical Models"

### *Supplementary Material*

#### 1 Supplementary Figures and Tables

##### 1.1 Supplementary Figures

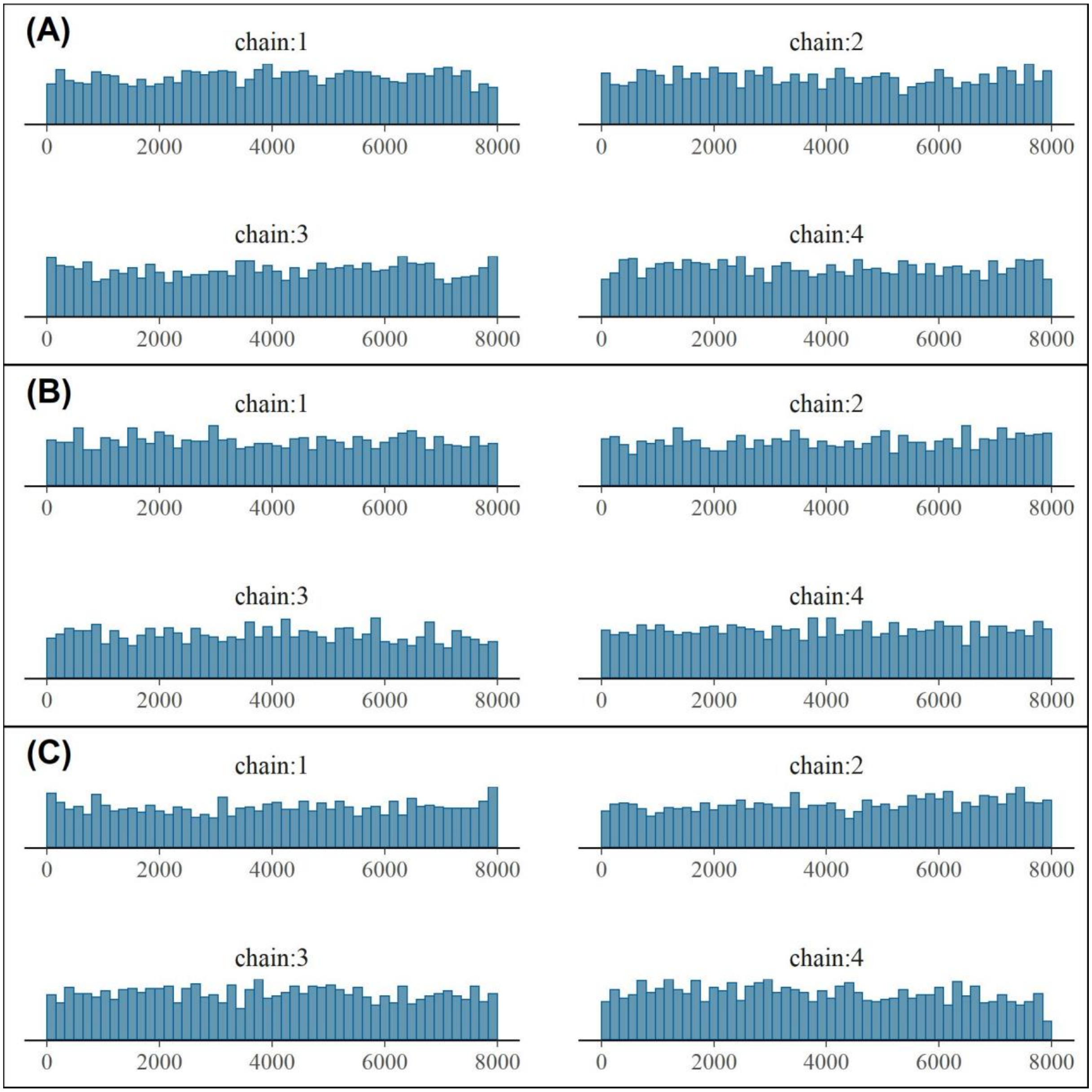


**Supplementary Figure 1.** Rank plots from posterior draws of the top-ranked occupancy model tracking the mixing of MCMC chains. Panels display rank plots for regression parameters $\{\alpha_l, \beta_l\}$ (**A**), parameters of the Gaussian process explaining temporal variation in detection probability $\{\sigma_{f_1}, \rho_{f_1}\}$ (**B**)

and spatial variation in occupancy $\{\sigma_{f_2}, \rho_{f_2}\}$ (**C**). For all panels, only parameters with the lowest tail-ESS (effective sample size) are displayed.

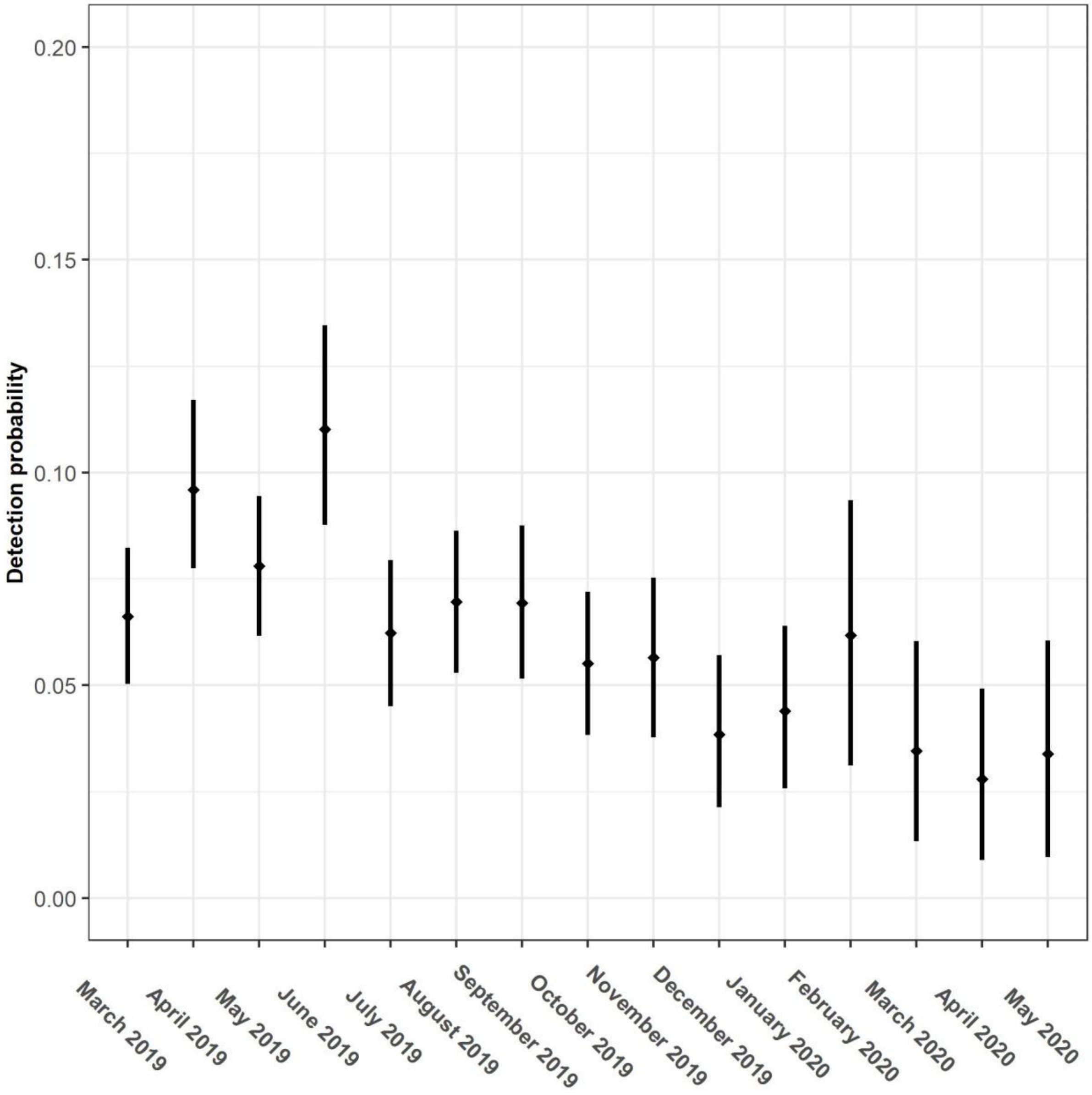


**Supplementary Figure 2:** Probability of detecting wild boar in response to the observation month. Posterior means (dots) and 95% highest posterior density intervals (vertical lines).

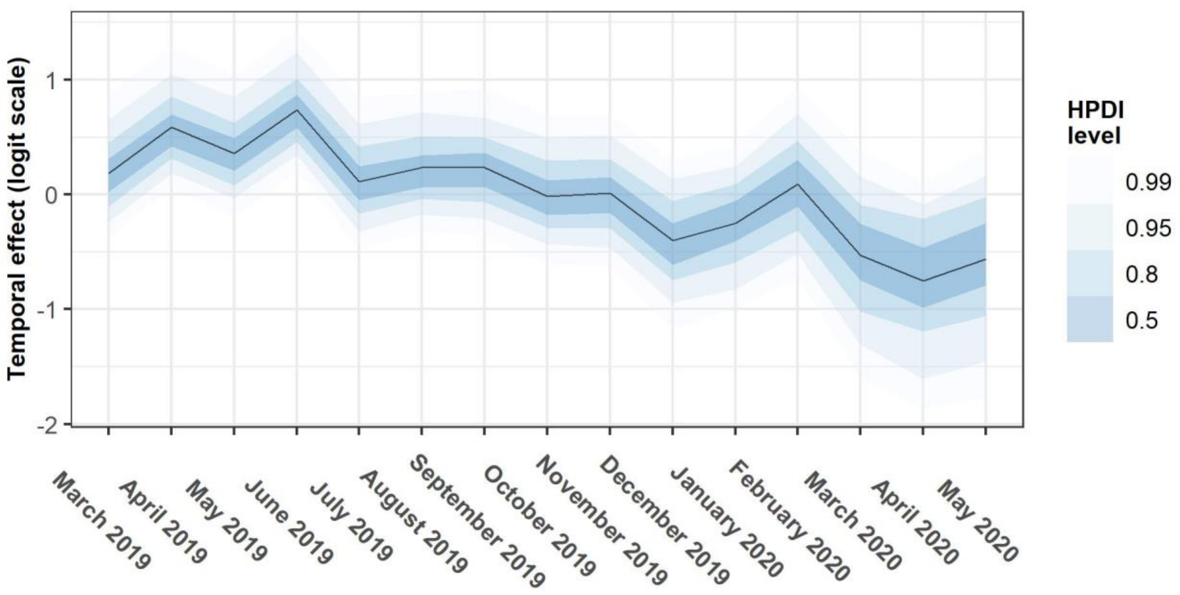


**Supplementary Figure 3:** Posterior mean and 50, 80, 90 and 95% highest posterior density intervals (HPDI) for the temporal effect (detection process) estimated by a Gaussian process. All values are presented on the logit scale.

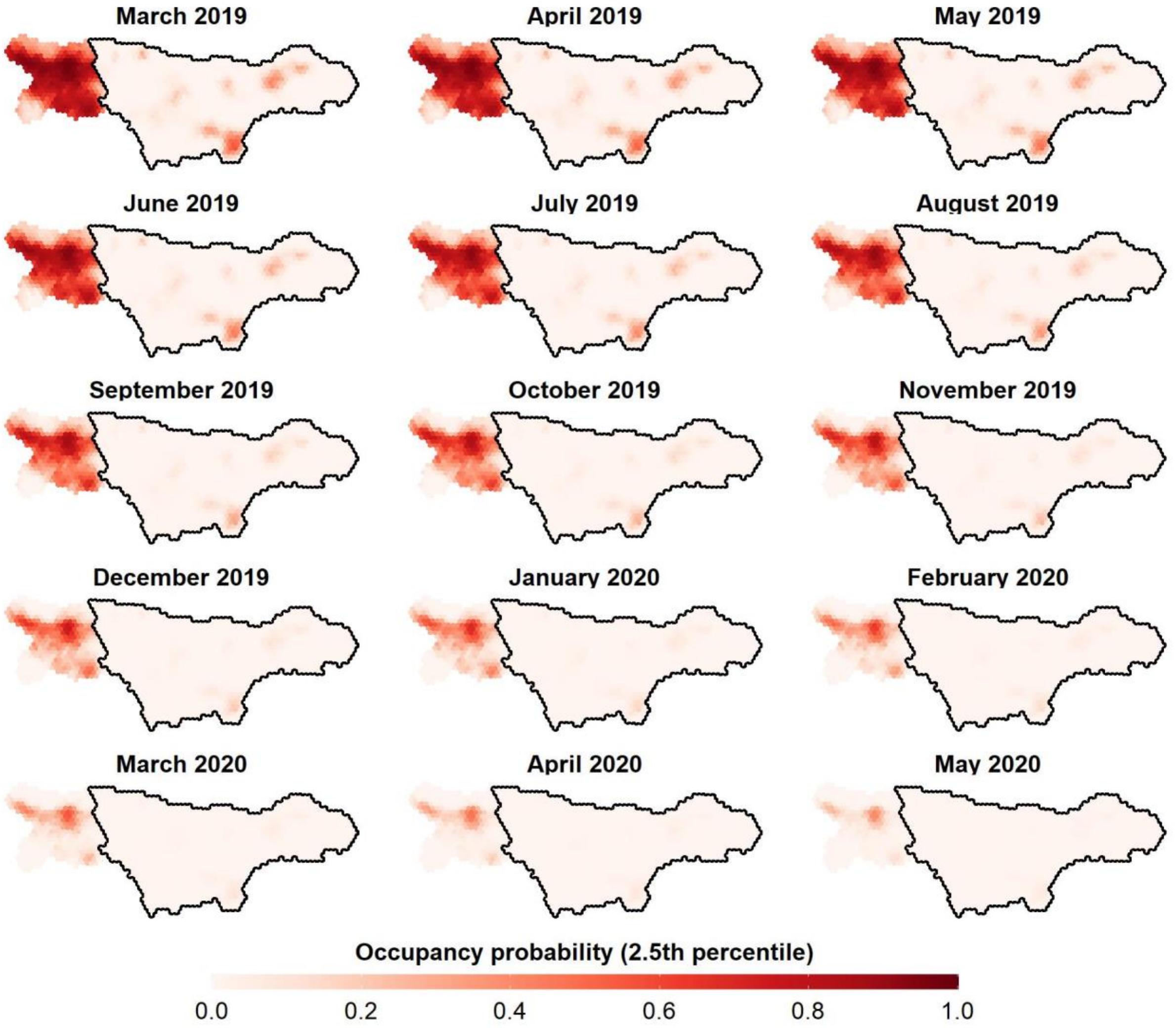


**Supplementary Figure 4:** 2.5th Percentile of posterior occupancy of wild boar in the ASF-infected (enclosed by the black line) and noninfected (non-enclosed) zone in the Wallonia (Belgium). Panels ranging from March 2019 until May 2020.

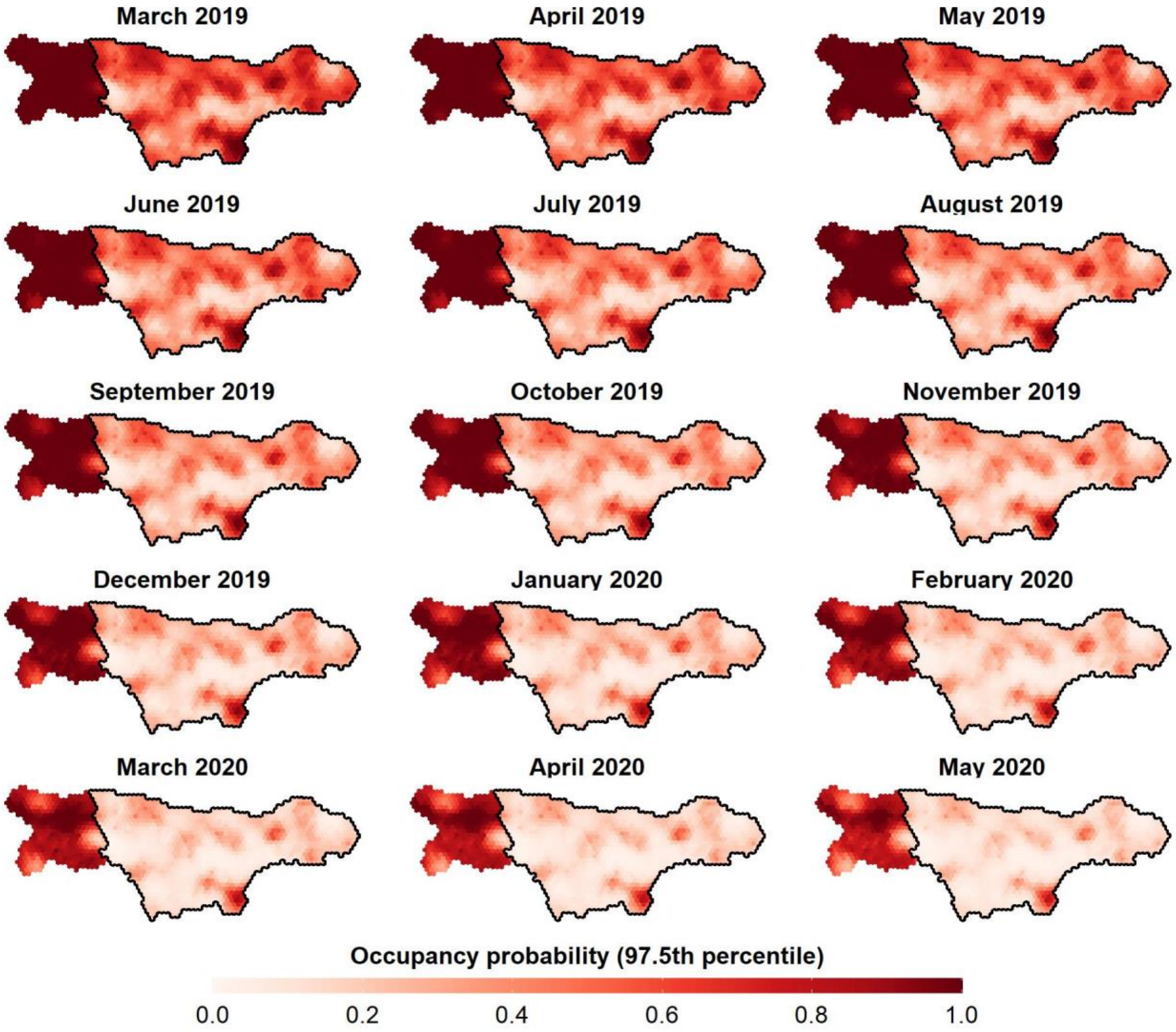


**Supplementary Figure 5:** 97.5$^{th}$ Percentile of posterior occupancy of wild boar in the ASF-infected (enclosed by the black line) and noninfected (non-enclosed) zone in Wallonia (Belgium). Panels ranging from March 2019 until May 2020.

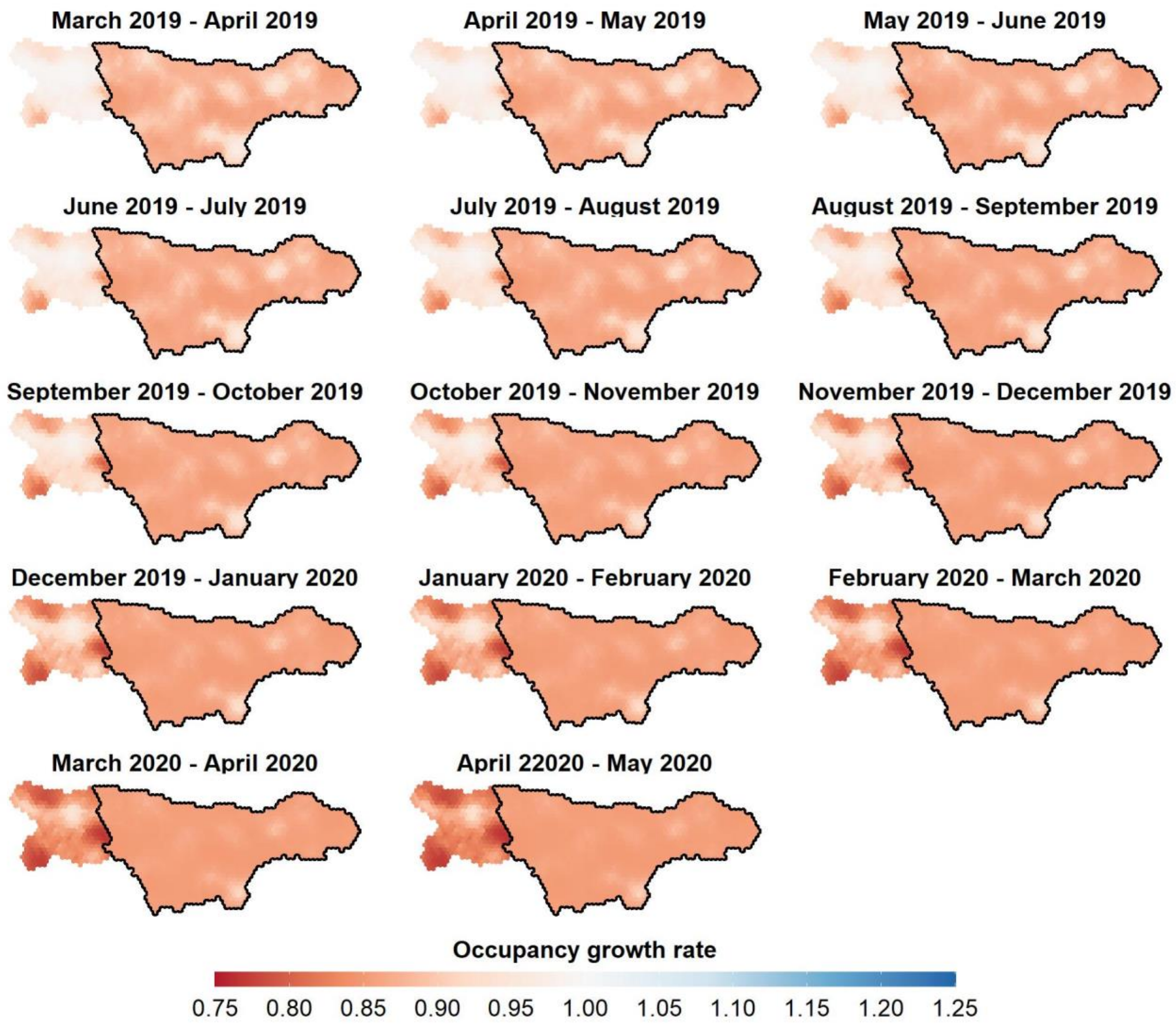


**Supplementary Figure 6:** Posterior mean occupancy growth rates of wild boar in the ASF-infected (enclosed by the black line) and noninfected (non-enclosed) zone in Wallonia (Belgium). Panels display growth rates derived from two consecutive months, ranging from March 2019 – April 2019 until April 2020 – May 2020.

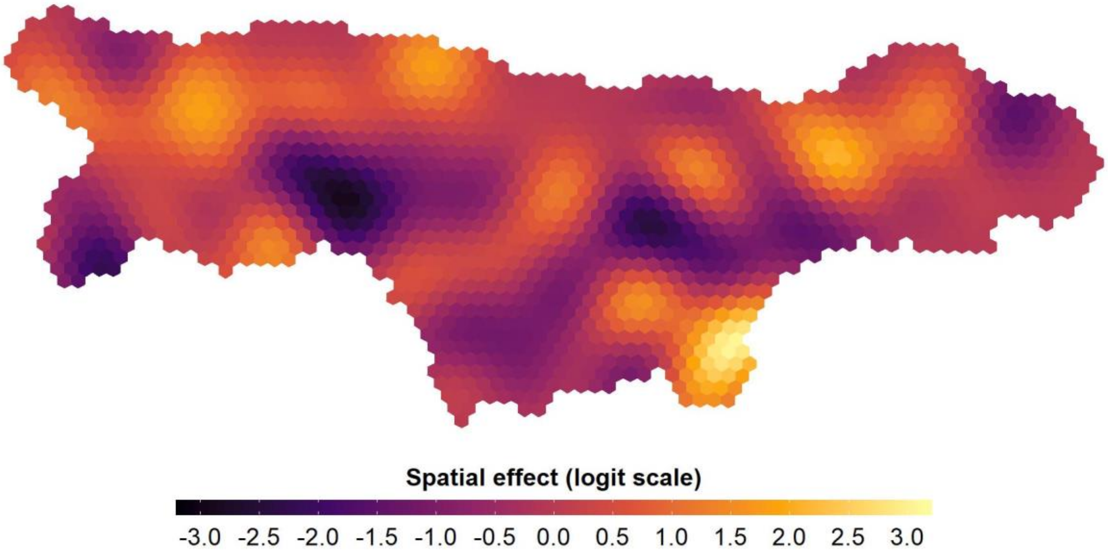


**Supplementary Figure 7:** Posterior mean of the spatial effect (occupancy process), estimated by a Hilbert-space Gaussian process. Mean values are presented on the logit scale.

#### 1.2 Supplementary Tables

**Supplementary Table 1:** Features and characteristics of the Snapshot Extra Black 12.0 l HD (Dörr) camera, as specified by the manufacturer.

| Camera feature | Characteristics |
|---|---|
| Motion sensor | Infrared |
| Trigger speed | 1/5 sec |
| Detection range | 14.6 m |
| Detection angle | 58° |

**Supplementary Table 2:** Sampling effort for each of the ASF management zones.

| Zone | Total Grids | Number Sampled | Sampled (%) |
|---|---|---|---|
| ASF-infected | 1135 | 69 | 6.08 |
| Noninfected | 348 | 23 | 6.61 |
| Excluded | 172 | 5 | 2.91 |

**Supplementary Table 3:** Predictions for *a priori* defined occupancy (step 1) and detection (step 2) models as defined in Table 1.

| Model | Prediction |
|---|---|
| Occupancy models (step 1) | |
| $\psi 1$ | P($\psi 1$): No selection. |
| $\psi 2$ | P($\psi 2$): Higher occupancy in noninfected sites. |
| $\psi 3$ | P($\psi 3$): Higher occupancy in noninfected sites with a high percentage of broad-leaved tree land cover class. |
| $\psi 4$ | P($\psi 4$): Similar as P($\psi 2$), additionally an overall linear (declining) trend in occupancy exists. |
| $\psi 5$ | P($\psi 5$): Similar as P($\psi 3$), but with different occupancy trends for ASF-infected and noninfected zones. |
| $\psi 6$ | P($\psi 6$): Similar as P($\psi 2$), with the temporal trend in occupancy best captured by a flexible (non-linear) process. |
| Detection models (step 2) | |
| $p1$ | P($p1$): No selection. |
| $p2$ | P($p2$): Lower detectability during spring and summer due to denser vegetation cover. |
| $p3$ | P($p3$): Lowest detectability during summer due to denser vegetation cover, followed by spring and autumn. Highest detectability during winter. |
| $p4$ | P($p4$): Temporal trend in detectability best captured by a flexible (non-linear) process. |

**Supplementary Table 4:** Odds ratios for posterior means and 95% highest posterior density intervals, extracted from the top-ranked model. Only regression parameters are reported.

| Parameter | Mean | 2.5% | 97.5% |
|---|---|---|---|
| $\alpha^{p}$ | 17.71 | 3.49 | 95.12 |
| $\beta^{\psi}_{ASF}$ | 0.01 | 0.00 | 0.08 |
| $\beta^{\psi}_{BL}$ | 1.48 | 0.97 | 2.35 |
| $\beta^{\psi}_{t}$ | 0.76 | 0.65 | 0.88 |
| $\beta^{\psi}_{ASF\cdot t}$ | 1.13 | 0.97 | 1.32 |
| $\alpha^{\psi}$ | 0.06 | 0.04 | 0.08 |

**Supplementary Table 5:** Posterior Mean and 95% highest posterior density values for zone-averaged occupancy ($\boldsymbol{\psi}_{t,z}$) estimates at observation month $\boldsymbol{t}$.

| $\boldsymbol{t}$ | **Month** | **Mean** | **2.5%** | **97.5%** | **Mean** | **2.5%** | **97.5%** |
|---|---|---|---|---|---|---|---|
| | | ASF-infected | | | Noninfected | | |
| 2019 | | | | | | | |
| 1 | March | 0.2352 | 0.0366 | 0.5399 | 0.8677 | 0.6342 | 0.9958 |
| 2 | April | 0.2131 | 0.0312 | 0.5015 | 0.8453 | 0.5954 | 0.9922 |
| 3 | May | 0.1923 | 0.0265 | 0.4638 | 0.8197 | 0.5544 | 0.9875 |
| 4 | June | 0.1730 | 0.0224 | 0.4269 | 0.7907 | 0.5110 | 0.9814 |
| 5 | July | 0.1550 | 0.0188 | 0.3911 | 0.7584 | 0.4656 | 0.9741 |
| 6 | August | 0.1385 | 0.0156 | 0.3568 | 0.7227 | 0.4177 | 0.9642 |
| 7 | September | 0.1234 | 0.0129 | 0.3243 | 0.6838 | 0.3686 | 0.9522 |
| 8 | October | 0.1097 | 0.0106 | 0.2937 | 0.6421 | 0.3187 | 0.9372 |
| 9 | November | 0.0972 | 0.0087 | 0.2651 | 0.5980 | 0.2691 | 0.9191 |
| 10 | December | 0.0860 | 0.0070 | 0.2386 | 0.5523 | 0.2209 | 0.8972 |
| 2020 | | | | | | | |
| 11 | January | 0.0759 | 0.0056 | 0.2142 | 0.5057 | 0.1747 | 0.8711 |
| 12 | February | 0.0668 | 0.0045 | 0.1917 | 0.4592 | 0.1339 | 0.8418 |
| 13 | March | 0.0588 | 0.0035 | 0.1712 | 0.4135 | 0.0984 | 0.8091 |
| 14 | April | 0.0517 | 0.0027 | 0.1527 | 0.3696 | 0.0698 | 0.7747 |
| 15 | May | 0.0453 | 0.0022 | 0.1360 | 0.3281 | 0.0476 | 0.7388 |

**Supplementary Table 6:** Numbers of wild boar culled per observation month ***t*** in each of the ASF management zones throughout the study period.

| ***t*** | **Month** | **Number of wild boar culled** | |
|---|---|---|---|
| | | ASF-infected | Noninfected |
| 2019 | | | |
| 1 | March | 56 | 53 |
| 2 | April | 23 | 55 |
| 3 | May | 23 | 23 |
| 4 | June | 9 | 72 |
| 5 | July | 3 | 100 |
| 6 | August | 9 | 52 |
| 7 | September | 17 | 12 |
| 8 | October | 8 | 39 |
| 9 | November | 24 | 36 |
| 10 | December | 18 | 37 |
| 2020 | | | |
| 11 | January | 10 | 11 |
| 12 | February | 18 | 17 |
| 13 | March | 14 | 15 |
| 14 | April | 5 | 10 |
| 15 | May | 11 | 2 |
